# Temperate bacteriophage induced in *Pseudomonas aeruginosa* biofilms can modulate bacteriophage and antibiotic resistance

**DOI:** 10.1101/2024.07.10.602839

**Authors:** Mark Grevsen Martinet, Bolaji John Samuel, Daniel Weiss, Mathias W. Pletz, Oliwia Makarewicz

**Affiliations:** Institute of Infectious Diseases and Infection Control, Jena University Hospital/Friedrich-Schiller University, Jena, Germany; Department of Orthopedics, Jena University Hospital, Eisenberg, Germany; Leibniz Center for Photonics in Infection Research, Jena University Hospital/Friedrich-Schiller University, Jena, Germany

## Abstract

Given the high levels of resistance in Gram-negative bacteria, phage therapy is garnering increasing attention. In Germany, a clinical study is already underway investigating a phage cocktail for the treatment of *Pseudomonas aeruginosa* in cystic fibrosis (CF) patients. In our study, we examined susceptibility to virulent phages and the PF1-like prophage and antimicrobial profiles and of *P. aeruginosa* isolates from a local cystic fibrosis cohort to identify correlations and lysogenic conversion of the prophegs. Consistent with other studies, prophage Pf4 is the most prevalent in this cohort and is activated in the absence of other influences during biofilm formation. These phages can be transferred to other strains that do not contain Pf1-like prophages, thereby influencing the dynamics of bacterial populations in the CF lung. This also rapidly leads to the emergence of a subpopulation resistant to the virulent phages, potentially complicating phage therapy. However, this subset also becomes more susceptible to most antibiotics commonly used in CF, which could be a useful treatment strategy. Interestingly, this bacterial subset lost its susceptibility to colistin, an important inhaled antibiotic in CF, which could lead to treatment failure. Our research highlights both the difficulties and potential strategies to improve treatment options for CF patients.

**Author summary:** We investigated 51 *P. aeruginosa* isolates obtained from CF patients for the presence of PF1-like prophages and characterized their susceptibility prior and after lysogenig conversion of the prophages to three virulent phages. Our study revealed that the temperate phage Pf4 is the most prominent PF1-like prophage in this cohort, undergoing lysogenic conversion during biofilm formation. The virions identified in the biofilm supernatants are superinfective and transferable to other prophage-free *P. aeruginosa* isolates, shaping population dynamics in the CF lung. Prophage reactivation results in the survival of a sub-population with reduced susceptibility to virulent phages, posing a potential challenge for phage therapy. However, this sub-population exhibited restored susceptibility to most CF-relevant antibiotics, presenting an intriguing therapeutic opportunity. Targeted prophage reactivation could sensitize multidrug-resistant *P. aeruginosa* in CF patients, enhancing or even restoring antimicrobial efficacy. Notably, this sub-population also showed a loss of colistin susceptibility, which may lead to treatment failure when colistin is used as an inhaled antipseudomonal antibiotic. Our findings highlight both significant challenges and potential therapeutic opportunities for optimizing the treatment of CF patients. However, these findings are preliminary and require further investigation, particularly regarding the possibility of lysogenic conversion in other prophages (such as *Siphoviridae*) and how these interplays with resistance to virulent phages. Further studies are ongoing and will enhance our understanding of the role of prophages in the pathophysiology of CF.

## Introduction

Cystic fibrosis (CF) is an autosomal recessive genetic disease caused by mutations in the cystic fibrosis transmembrane conductance regulator (CFTR) gene. CF affects millions of people worldwide and manifests primarily by production of thick, sticky mucus in the airways leading to life-threatening respiratory complications and infections. The CF mucus provides an ideal niche for pathogens to colonize, evade immune defences, and persistently infect the lungs (1). In particular, infections with *Pseudomonas aeruginosa*, a highly adaptable and virulent bacterium, are associated with strongly increased mortality in CF patients (2). Therefore, a key focus of CF therapy is the prevention of PA infections (3). If there is positive evidence of *P. aeruginosa*, aggressive therapy with various antibiotics at the same time, oral and inhaled, is used, and this is not always successful either. A significant factor contributing to the high resistance of *P*. *aeruginosa* to therapies is its ability to form biofilms. These biofilms protect the embedded bacteria from external influences such as antibiotics and the immune system. Additionally, the bacteria transition into metabolically inactive forms, allowing them to persist for extended periods and cause recurrent infections(4,5). According to the Cystic Fibrosis Foundation (https://www.cff.org), approximately 60% of CF adults suffer from infections or chronic colonisation of the lung by *P. aeruginosa*. Therefore, alternative therapies that address more specific *P. aeruginosa* and their biofilms are urgently needed.

Bacteriophages (thereafter named phages) are viruses that infect and kill bacterial cells with high specificity. Virulent phages replicate immediately upon infecting bacteria, leading to their lysis and are considered as potent therapeutics against difficult-to-treat infections antibiotic-resistant bacterial. Phage therapy offers the prospect of precision medicine, where tailored phage cocktails can be deployed to combat specific bacterial strains within individual CF patients (6). Phage therapy in CF is already being explored in clinical trials. According to the ClinicalTrials.gov, five trials in CF-patients with phage-cocktails against *P. aeruginosa* are registered (as December 2023), with two phase 1/phase 2 trials being completed (NTC04596319 on AP-PA02, and NCT04684641 on YPT-01, both study results are still unpublished), an recruiting follow-up phase 2 trial on AP-PA02 (NTC05616221), another recruiting phase 1/phase 2 trial on WRAIR-PAM-CF1 (NCT05453578), and one active but currently non-recruiting trial on BX004-A (NTC05010577). This indicates the upcoming interest on phages as potent antimicrobials.

Temperate phages, integrate into the bacterial genome as prophages, remaining there until reactivation (lysogenic conversion). Those are known to contribute to horizontal gene transfer (7) and are thus less potent as antimicrobial agents. Prophages are common in bacteria, particularly in *P. aeruginosa*. They can switch spontaneously or after induction by various intracellular (e.g. DNA damage) or external signals (e.g., foods and drugs) to the lytic cycle to form and release new phage particles (8). This is known as lysogenic conversion. However, interaction between therapeutically used virulent phages and prophages is still not well understood.

Prophages are increasingly recognized as an important component in the complex interplay between host, pathogen and environment associated with CF (9–11). Prophages can carry genes that confer a range of benefits to their host bacterium, including antibiotic resistance, virulence factors, and metabolic factors for adaptation to environmental stress (12). For instance, prophages can encode toxins and secretion systems that enhance the bacterium’s pathogenic potential, contributing to tissue damage and exacerbation of lung infections (13). These genetic payloads can profoundly impact *P. aeruginosa’s* ability to colonize, persist, and resist immune clearance within the CF lung. Furthermore, prophage induction can lead to the formation of biofilms, a critical factor in the chronicity of *P. aeruginosa* infections in CF-lung (14).

This dynamic interaction between prophages and the host bacterium shapes the clinical course of CF pathogenesis, posing significant challenges to the management and treatment of this complex disease, also in the view of phage therapy. Therefore, we aimed to analyse the prophage profiles of *P. aeruginosa* strains from a German CF cohort and investigate their ability of lysogenic conversion and its impact on antibiotic treatment and treatment with virulent phages.

## Results and Discussion

### Demographics of the CF cohort

A total of 51 PA isolates derived from 49 specimen of 29 CF patients were examined retrospectively in this study. From 12 of these patients, multiple isolates were obtained and examined: In six cases, two phenotypically different *P. aeruginosa* isolates were derived from a single sample, and in two cases, three different isolates were identified from one sample. The remaining isolates from these patients were collected during different visits. For the other 17 patients, only one isolate was analysed, even though *P. aeruginosa* was isolated from them at other visits. Due to the small sample size and the number of repeated isolates from the same patients, we do not consider a demographic analysis of the patients to be useful for the purpose of the project. However, factors such as age and gender could play a role, which is why these were recorded descriptively for each individual isolate. The range of the patients’ age ranged from 7 to 57 years, with some more frequent samples at age range between 32 and 35 and an average age of 30 (Supplementary Material Figure S2). The distribution of sex was almost equal with 45% male and 55% female patients. The specimen distribution was 41% sputum (n = 20), 27% throat swap (n = 13) and 14% nasal lavage (n = 7). Mucoid growth on agar plates was observed in 49% (n = 24) of the isolates.

Correlated to the number of specimen (n = 41), additional other species were found in 61.0% the samples, with *S. aureus* being most prominent (29.3%), followed by not further differentiated regular mouth flora (24.4%). *Escherichia coli*, *Candida albicans* and *Aspergillus fumigatus* were found only one time each and accompany the *P. aeruginosa* isolates used in this study.

### Resistances of the *P. aeruginosa* strains

For an initial detection of *P. aeruginosa* with pulmonary exacerbation, intravenous (iv) antibiotic therapy is recommended. Various substances can be used, typically depending on the resistance profile of the isolates. To improve the success of eradication, therapy is usually followed by inhalation of tobramycin or colistin, along with oral administration of ciprofloxacin (15). Other inhaled antibiotics currently available include aztreonam lysine and levofloxacin. These can also be used as needed.

Resistance profiles of the 51 isolates were determined during routine diagnostics as minimal inhibitory concentration (MIC) and interpreted according to the EUCAST guideline as susceptible, intermediate and resistant (Supplementary Material Figure S3 A). All isolates were tested against ceftazidime (CAZ), piperacillin/tazobactam (PIP/TAZ), meropenem (MER), ciprofloxacin (CIP) and tobramycin (TOB, here for one patient TOB was not determined for unknown reason). These antibiotics constitutes the essential anti-pseudomonal therapy in Germany (15). Only 2% and 4% of the isolates were susceptible to the β-Lactam antibiotics CAZ and PIP/TAZ respectively, resistance was present in 33% and 35%. Most of the isolates showed intermediate MICs to these antibiotics. 53% of the isolates were susceptible and 14% were resistant to MER, a carbapenem and last-resort β-lactam. Intermediate MIC_MER_ were found in 33% of the isolates. Resistance to CIP was present in 65% and susceptibility in 10% of the isolates, while 25% of the isolates showed intermediate MIC_CIP_. The MIC for tobramycin (TOB), most used as inhaled therapy, was tested in 50 strains. 56% of the isolates were susceptible and 34% were resistant to TOB. Intermediate MIC_TOB_ were found in 10% of the isolates.

Additional antibiotics were tested against some strains that are used for inhaled treatment in chronic *P. aeruginosa* infections (Supplementary Material Figure S3 B). Colistin (COL) is also commonly used as inhaled therapy, but the MIC_COL_ was assessed in only 28 isolates and resistance was found in 13% of cases. The reason for the infrequent testing of MIC_COL_ is the empirical efficacy of inhaled therapy despite high MIC values, allowing the antibiotic to be successfully used in CF therapy without strict adherence to the MIC. This may be due to the high local antimicrobial concentration achieved through inhaled application. This effect is also observed with levofloxacin (LEV) and aztreonam (ATZ), which are alternatives to TOB or COL. LEV and ATZ were tested in only 30 and 18 isolates, showing resistance rates of 77% and 22%, respectively.

Amikacin (AMI) is another antimicrobial that can be administered intravenously or inhaled to prevent or treat bacterial infections in CF patients. The MIC_AMI_ was tested in 46 isolates, with susceptibility observed in 61% of the isolates. However, AMI is rarely prescribed in the current cohort and is primarily used for non-pseudomonal infections, such as *Mycobacterium abscessus*.

The antimicrobial susceptibility showed in general positive correlations between the compounds (Supplementary Material Figure S3) that corresponds to the often observed multidrug-resistance in CF-isolates (16)Interestingly there was a weak but negative correlation between COL and ATZ susceptibility that might result from the low number of isolates with assessed MIC_ATZ_. The antibiotic therapy in CF is guided by observed efficacy and tolerability. Since inhaled formulations can be administered locally at higher doses, MIC values play a less significant role, and this negative correlation has no practical relevance.

### Distribution of prophage Pf1-like specific genetic element

The Pf phages belong to the filamentous inoviruses infecting the bacteria via type IV pili (17). Different Pf1-like genetic elements can be found in *P. aeruginosa*. In a prior PCR-screening study from 2015, out of 241 analysed *P. aeruginosa* strains, 60% contained at least 1 genetic element of the Pf-1-like family with Pf4 being most prevalent (18).

Using the previously described universal primers PfUa (19) and PfUb (18), we assessed the presence of Pf1-like prophage signatures in the CF-isolates via PCR (Figure 1, A). In total, 59% and 65% of the isolates were positive for PfUa and PfUb amplicons, respectively, with 55% being positive for both PCR products. The PfUa and PfUb frequencies in the CF cohort were higher compared to the analyses of Mooji *et al*. and Knezevic *et al*., where PfUa amplicons were identified in 44% and 41%, respectively, and PfUb was identified in 52% of *P. aueruginosa* strains from different sources including environmental strains (18,19).

**Figure 1.**
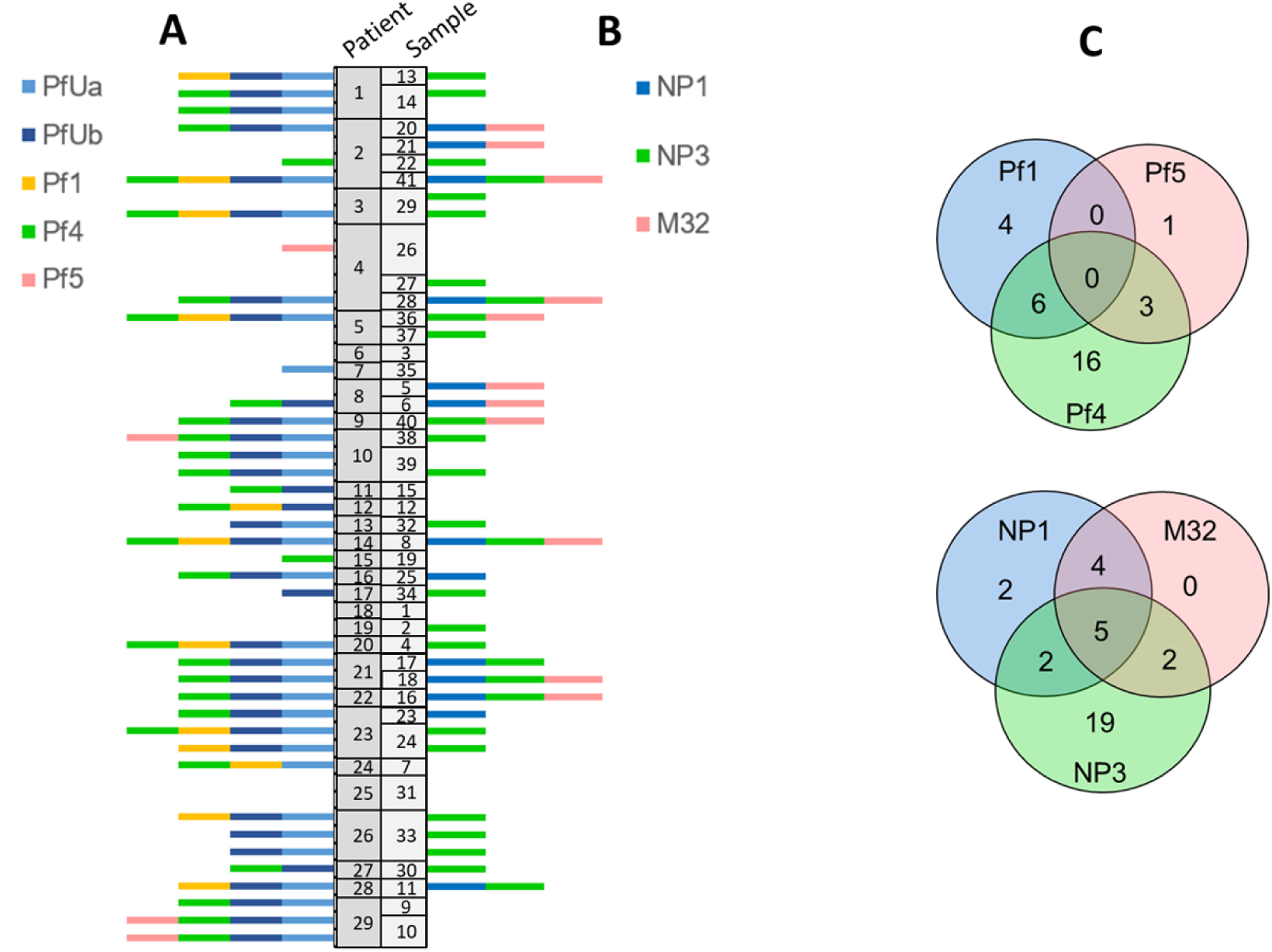
Distribution of the prophage-related genetic elements determined by universal (PfUa and PfUb) primers or the specific primers for Pf1, Pf4 and Pf5 (A) and the susceptibility to virulent phages NP1, NP3 and M32 (B). The Y-axis indicates the individual patients ID (given randomly) and samples IDs and the X-axis indicates presence of the prophage amplicon (A) or susceptibility to virulent phage (B) as indicated in the legend (respective colour). (C) Venn diagrams of the distributions of the prophage elements and susceptibility to the virulent phages within the CF isolates.

We used primers that have been previously designed and proved to bind at specific genetic elements related with subgroups of the Pf1-prophages, namely Pf1, Pf4 and Pf5 (18,19). The most abundant specific amplicon was for Pf4-prophage with 55% followed by Pf1 in 24% of the isolates. Comparing to the previous work of Knezevic *et al.* (18), Pf1 (17.8%) prevalence was only slightly higher in our CF cohort and simultaneous presence of Pf4 with the general PfU and Pf1 amplicons did not significantly differ, but the occurrence of the Pfu4 amplicon in our cohort was more than twice as high (Figure 1 A). This explains also the higher frequency of the PfU amplicons in the CF cohort and indicates that there might be a predisposition for specific *P. aeruginosa* subtypes in CF patients. Pf1 phage is the only one known extrachromosomally persisting prophage of the Pf1-like (20), while Pf4 phage has been related to virulence of *P. aeruginosa* in a mice model (21).

In accordance with Knezevic *et al*. (18), the Pf5 amplicon was only identified in 4 isolates (7.8% vs 6.6%) and specific signatures of all three prophages were not identified in the CF isolates (Figure 1 C).

In our cohort, only 7.8% of the isolates (n = 4) were solely positive for the universal primers and in 29% (N = 15), none of these two amplicons were identified and in 24% (N = 12) no other specific prophage fragments were amplified (in comparison: 39.8% in the work of Knezevic *et al*.). Both signatures correlated well with each other but showed only low positive correlation to the specific phage amplicons of Pf1 and Pf4, and almost no correlation to Pf5 (Pearsons’s rank (r_p_) =0.71, Supplementary Material Figure S3). This indicated that the universal primes might fail to cover sufficiently the Pf1-prophages in the CF cohort. The missing amplicons of the PfU-primers might also indicate the presence of Pf7 phage clade as previously estimated (18) or simply result from genetic alterations in the CF-isolates.

A recent study in a Standfort and Danish cohort revealed that the isolates from older CF-patients were more likely to be Pf-positive (22). However, in our cohort, there was only a weak positive trend to Pf1 amplicon and age.

### Susceptibility to the virulent phages NP1, NP3 and M32

The CF-isolates were investigated for their susceptibility to the three virulent phages NP1, NP3 and M32, which exhibit broad host ranges against *P. aeruginosa* strains (23,24) being potential for phage therapy also to treat CF patients. Therefore, we aimed to evaluate the susceptibility of the strains in the CF cohort against those phages (Figure 1 B). We differentiated between susceptibility, reduced susceptibility and resistance.

Susceptibility to NP1 was determined in 25%, and in 55% to NP3 and in 22% to M32 of the isolates; while resistance was present in 67% to NP1, 35% to NP3 and 73% to M32. The residual fractions showed reduced susceptibility on the agar plates visible as isolated colony growth. As reduced susceptibility might result in a treatment failure in phage therapy, we suggest interpreting this phenotype rather as resistance for diagnostic purpose.

Susceptibility to all three virulent phages was determined in 33% (N = 17) of the CF-isolates, but 10% of the strains showed reduced susceptibility or resistance to all three virulent phages. 37% were susceptible only to NP3 and 3.9% only to NP1, while no of *the P. aeruginosa* strains showed susceptibility solely to M32 (Figure 1 C). This indicates a general high resistance of the CF *P. aeruginosa* strains against these virulent phages that might result in treatment failure when applying phage therapy.

Despite different phage classes, the resistance to NP1 and M32 correlated positively to each other (Supplementary Material Figure S4) suggesting identical receptors for invasion of both phages. It might be interested to analyse if these phages also compete when used together in a cocktail resulting in antagonism and reduced affectivity (25).

### Correlation between prophages, and susceptibilities of phages and antimicrobials

There were no notable correlations between resistance to the virulent phages and the presence of the prophages, only some trends might be estimated. However, interpretation of the data is difficult as there are hardly any comparable analyses in other descriptive studies.

A negative trend of the r_P_ between Pf1 and NP3 was present (Supplementary Material Figure S4). Interestingly, the presence of Pf5 elements showed low but positive r_P_ values. It might be interesting to investigate more isolates as the prevalence of Pf1 and Pf5 elements was low, to figure out if presence of Pf1 elements might be a suitable indicator for increased susceptibility and if Pf5 is an indicator of reduced susceptibility specific virulent phages used therapeutically.

There is a weak negative trend between the presence of the Pf1 and resistance to MER. Positive trends were found between TOB resistance and Pf5 presence and NP1 resistance and TOB or CAZ resistance. ATZ resistance showed also positive trend to the presence of all prophages tested and a positive correlation to the resistance against the virulent phages. However, we assume that any correlation of ATZ-susceptibility was biased due to the not frequently assessed MIC_ATZ_.

### Clonality of the isolates

It is noticeable that the isolates obtained from one patient either at different time points, or in the same specimen differ greatly in their properties. Even if the prophage pattern was often similar, at least for isolates from a single specimen, the susceptibility to virulent phages and the resistance pattern were generally different. This highlights the well-documented variability in *P. aeruginosa* strains that occupy the CF lung. This variability underscores the complexity of managing infections in CF patients, as the differences in phage susceptibility and antibiotic resistance can significantly impact treatment efficacy and outcomes (26).

### Prophage induction in *P. aeruginosa* biofilms

Hypoxia, nutrient limitations, and host immune responses can all stimulate prophage induction, leading to bursts of phage production and bacterial cell lysis(27). All these conditions also apply to biofilms in the CF lung. Therefore, we studied the reactivation of the prophage Pf4 as most abundant one during formation and maturation of the biofilms in the laboratory strain PAO1 and two clinical isolates (CF-PA75 and CF-PA83). These strains were selected because they all contain the Pf4 signature, and they were susceptible to all three exogenous phages. The clinical isolate CF-PA83 was also positive for Pf1, while PAO1 and CF-PA75 were both Pf1-negative. Additionally, isolate CF-PA6, which does not contain any of the prophage signatures was used as negative control.

Presence of phages with lytic activity in biofilm supernatants was assessed daily by the spot tests against the host strain (self-infectivity) and the presence of reactivated phages was confirmed by PCR for the supernatants after 5 days of growth (Supplementary Material, Figure S5). Pf1 could not be terminated in the supernatants of any of the strains, while all supernatants except of CF-PA6 were positive for Pf4 and Pf5 was identified in the supernatant of CF-PA83.

We assessed the phage titre in the biofilm supernatants in a time dependent manner during the biofilm maturation (Figure 2 A) in PAO1. In the laboratory strains PAO1, the Pf4 titre increased strongly in a time dependent manner achieving a maximum after 5 days and slightly doped down at day six. The supernatants of the clinical isolates showed already a high phage titre after one day of incubation that also increased by two log-magnitudes with the time. The differences in the titres between the supernatants obtained from clinical isolates and the laboratory strain might result from the generally higher cross-infectivity and a time-dependent increase in the self-infectivity in PAO1. However, this phenomenon was not investigated in detail in this study. The high cross-activity of the supernatants from biofilms of clinical isolates in PAO1 already after one day of biofilm growth indicates that prophages can be easily reactivated and released by CF-isolates. As Pf4 was demonstrated to impact virulence of *P. aeruginosa* (21), the lysogenic conversion can potentially worsen the lung condition of CF patients.

**Figure 2.**
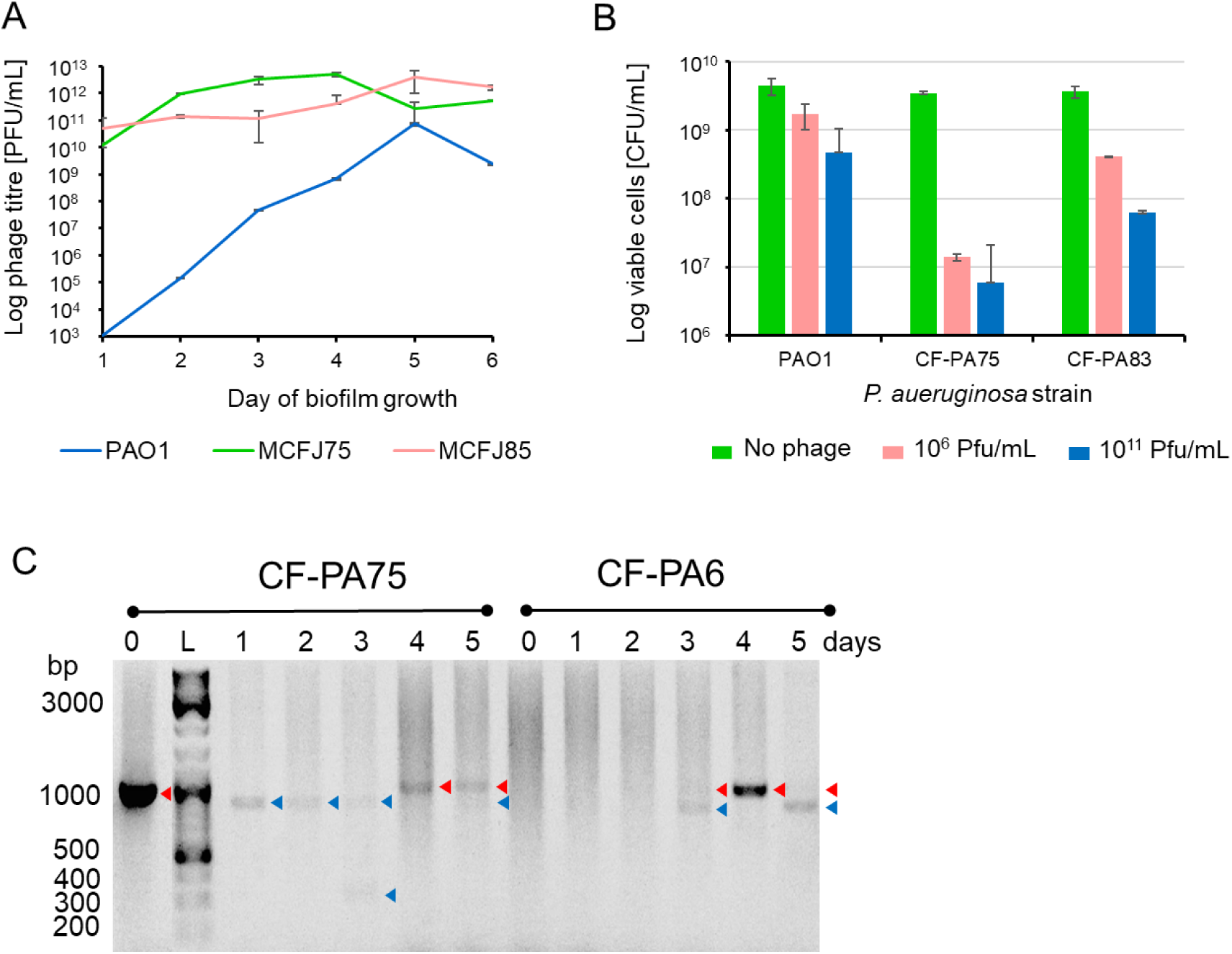
Pf4 induction and infectivity in biofilms of selected clinical isolates and PAO1. A) Lytic activity of the supernatant across several days of biofilm growth. The phage titre (PFU/mL) was assessed in PAO1 for all supernatants (see legend). B) Self-infectivity of the induced temperate phages on 24 h old biofilms of PAO1, CF-PA75 and CF-PA83 determined as viable bacteria (CFU/mL) after 16-18 h of phage treatment. Two different phage concentrations (10^6^ and 10^11^ PFU/mL) were tested. C) Transmission of Pf4 between CF-PA75 and CF-PA6 (Pf4-negative) in a mixed biofilm. The biofilms were growth between 1 to 5 days. The PCR was performed from 5 randomly selected colonies grown after biofilm resuspension on selective agar plates. 0 = CF-PA75 and CF-PA6 ancestor strain, L = molecular ladder (as indicated left hand). Red arrows indicate the expected 1001 bp band, blue arrows indicate a smaller PCR fragment of approximately

We further tested the self-infectivity on 24-h old biofilms threatening the corresponding strains overnight (16-18 h) with two different concentrations of the phage lysates (10^6^ and 10^11^ PFU/mL, obtained from the PEG enrichment). The viable bacteria were assessed after washing and resuspending of the biofilms in fresh LB-broth as CFU/mL on LB agar (Figure 2 B). The phage lysates reduced the bacteria in a concentration dependent but not linear manner. On PAO1 biofilms, the Pf4 virulence was lower compared to the clinical isolates and 10^11^ PFU/mL resulted in only one log-reduction. In CF-PA75, the Pf4 virulence was highest with 2-log reduction by 10^6^ PFU/mL, however, the 10^11^ PFU/mL only resulted in an additional one log reduction. In CF-PA83 the CFU/mL were reduced stepwise by one log each concentration. Here we cannot strictly separate if the effect corresponds to Pf4 or Pf5, but the lower effectivity might result from competitive effects of both phages during infection as both recognise the type IV pilus.

Presence of prophages usually results in immunity to infections by related phages (28). The self-infection suggests the ability of Pf4 (and possibly Pf5) to superinfection. The observed nonlinear correlation between PFU/mL and CFU/mL could be an interplay between the superinfectivity of the temperate reactivated phages against its host and the development of phage resistance mechanisms in biofilms. The superinfection in turn indicates genetic alteration of the Pf4 and Pf5 phages during the reactivation process to overcome bacterial immunity. Superinfective induced prophages have been shown to exhibit a higher frequency of mutations, which can result in the irreversible switch of prophages from a non-lytic phenotype (such as the filamentous Pf4 phage) to a lytic phenotype(27).

It must be noted that the high concentration of phages used for reinfection, which was almost 2 logs higher than the initial concentration of bacteria did not result in elimination of the bacteria. The results showed that approximately 10% (PAO1), 1% (CF-PA83), or 0.1% (CF-PA75) of the biofilm populations were not susceptible to the reactivated phage. This means that this subpopulation would prevail over time. It remains unclear whether this resistance is due to a newly acquired resistance mechanism of the host or of the integrated prophage restoring the immunity to reinfection. We will address these questions in in further studies.

### Transmission of Pf4

To speculate the possibility of transfer of prophages between the *P. aeruginosa* population in CF patient, the transmission of the Pf4 prophage within a mixed biofilm was investigated. CF-PA75 (meropenem resistant) contains the Pf4 prophage genetic element, while the CF-PA6 (tobramycin resistant) does not. Both isolates were mixed equally and to form mixed biofilms. The biofilms were resuspended, and the embedded viable bacteria were selectively grown on agar plates daily. From five randomly selected and mixed colonies of each plate, the Pf4-specific fragments were amplified (Figure 2C).

For CF-PA73, a slightly smaller PCR fragment was detected in the colonies collected during the first three days of biofilm formation. This fragment was also present at a lower level after five days. At day three, also a 300 bp amplicon was visible. The expected amplicon of 1001 bp was observed on the fourth and fifth day. Similar-sized amplicons were observed in CF-PA6 after three days of mixed biofilms clearly indicating a transmission of Pf4 between the strains. The difference in amplicon size suggests genetic alterations in the Pf4 capsid gene, which might correspond to Pf4 resistance in the surviving subpopulation. However, the amplicon was not sequenced thus we cannot comment on the changes. This observation is worth pursuing further, which we are investigating in currently ongoing studies.

### Resistance to virulent phages after prophage induction

Superinfection can drive the evolution of bacteria and can lead to increased genetic diversity resulting in resistance to other phages (29). Therefore, we further elucidate if Pf4 (*Inovirdae*) reactivation affects susceptibility of the bacteria to the non-related virulent phages NP1 (*Siphoviridae*) and NP3 (*Myoviridae*) and M32 (*Podoviridae*).

Biofilms grown from 1 to 5 days were resuspended in fresh LB broth after removal of the supernatants. After serial dilution, the bacteria were grown on agar plates. Randomly selected colonies (in total 32) were subjected to the cross-streak agar assay using the virulent phages NP1, NP3 and M32 to assess the fraction of susceptible and resistant clones. We also considered those clones where sensitivity was reduced due to individual colonies growing beyond the viral inoculation line (Figure 3).

**Figure 3.**
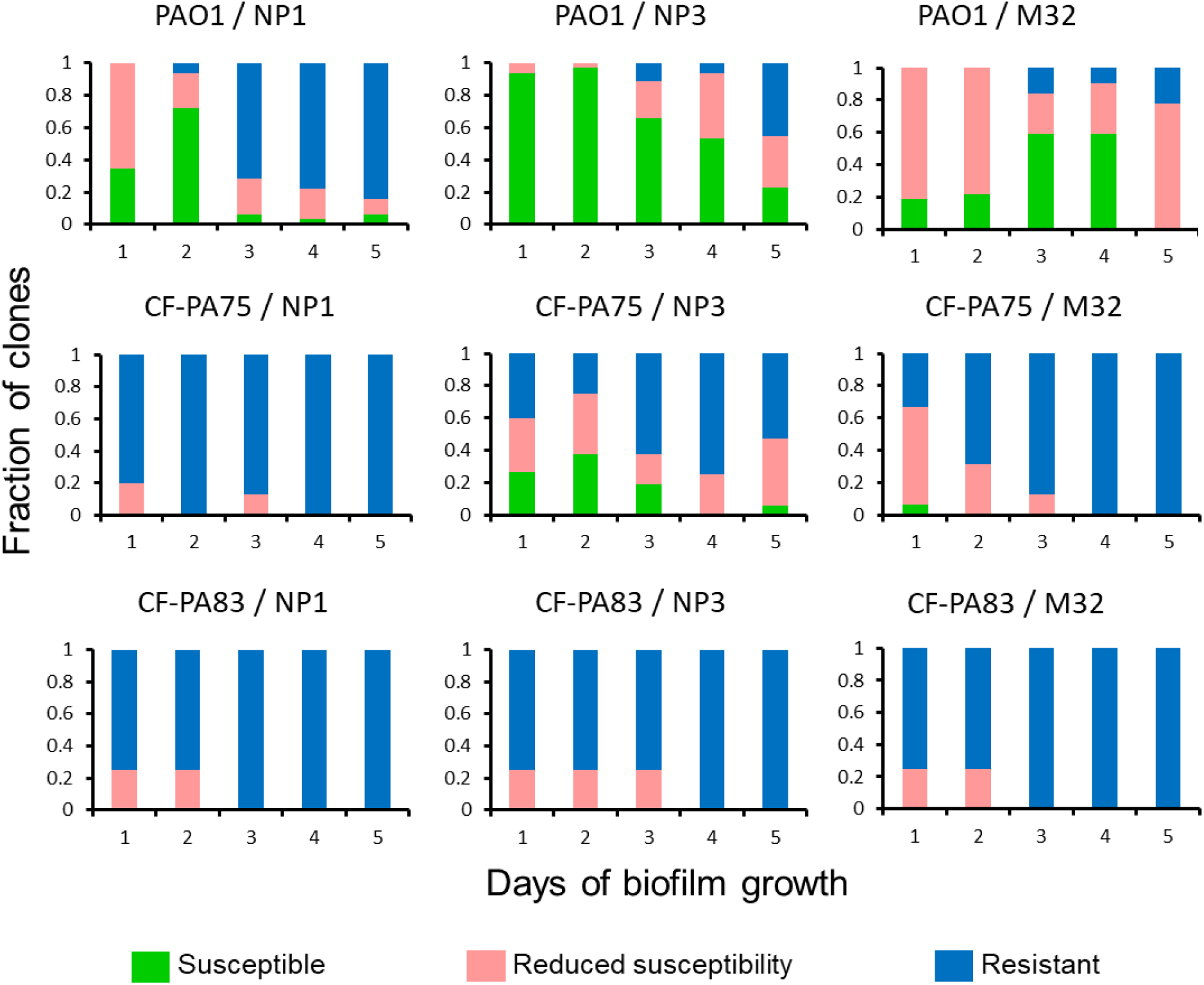
Development of resistance to virulent phages in selected clinical isolates and PAO1. Resistance profiles to virulent phages NP1, NP3 and M32 were assessed for biofilm embedded clones (daily N= 32 colonies randomly selected) in time dependent manner.

All bacterial strains (PA01, CF-PA75 and CF-PA83) were tested susceptible to the three virulent exogenous phages before biofilm formation. After one day of biofilm formation, there was already high fraction of PAO1-clones with reduced susceptibility to NP1 and M32 and high fractions of resistance clones to all phages in the clinical isolates. The biofilms of the laboratory strain PAO1 showed generally the highest fractions of susceptibility to the virulent phages, in particular to NP3. For NP3 susceptible fractions were also observed in CF-PA75 clones up to 5-day old biofilms. The increase in resistant fractions in the biofilms was observed in a time-dependent manner in all strains and against all phages. However, this development was particularly rapid in the clinical isolates. After three or four days, most clones showed resistance the virulent phages, particularly to NP1 and M32.

### Changes in antibiotics resistance profiles after prophage induction

Changes in antibiotic susceptibility during biofilm growth over 5 days were monitored daily. Each day, 32 colonies grown from the resuspended biofilms of PAO1 and CF-PA75 were randomly subjected to antimicrobial susceptibility testing (AST). The antimicrobial classes and their representative drugs used in this analysis included cephalosporins (ceftazidime and cefepime), carbapenems (imipenem and meropenem), broad-spectrum penicillins combined with β-lactamase inhibitors (piperacillin-tazobactam), fluoroquinolones (ciprofloxacin and levofloxacin), aminoglycosides (gentamicin, tobramycin, and amikacin), and polymyxins (colistin) (Figure 4).

**Figure 4.**
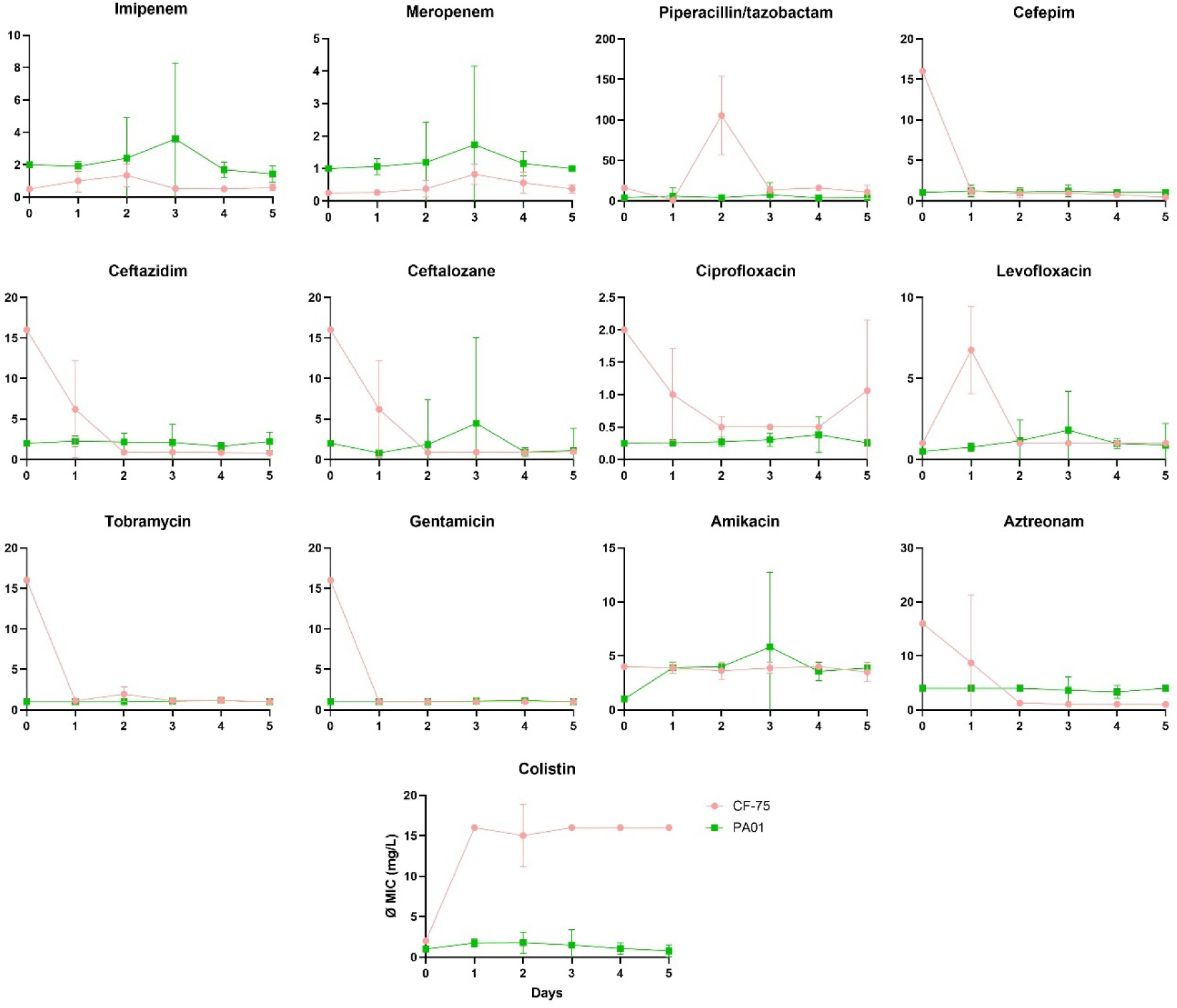
The effect of lysogenic conversion of the temperate bacteriophages on the susceptibility towards antibiotics. The y-axes represent the minimal inhibitory concentration (MIC) of the corresponding antimicrobial (as indicated above each graph). X-axes indicate the growth of the biofilm. The MICs were assessed for 32 clones for every day of biofilm growth and shown as mean and standard deviation

The origin bacterial strain PAO1 strain (time point 0) was sensitive to all antibiotics. The clones obtained from biofilms showed increase MICs for β-lactams and fluroquinolones in selected colonies that interestingly is most pronounced at day 3 of biofilm growth. This was also the time frame, where first phage transmissions were detectible in the previously prophage-free recipient in mixed biofilms. Therefore, we assume that the observed elevation of the MICs in PAO1 might correlate with the prophage induction and superinfection. However, the MICs changes were below the resistance breakpoints.

The CF-PA75 original strain showed MICs above the clinical breakpoint (resistance) for several antibiotics:128 mg/L for piperacillin, 16 mg/L each for ceftazidime, ceftalozane, cefepime, gentamicin, tobramycin, and aztreonam, 2 mg/L ciprofloxacin. Similarly to PAO1, the MICs of carbapenems increased during biofilm formation and reached the highest values after three days - with values significantly above the clinical breakpoints for imipenem (4 mg/L) in selected colonies on third day of biofilm growth. The MIC values for cephalosporines, aztreonam, ciprofloxacin, and aminoglycosides (except for amikacin that showed low MIC values over the whole time), declined with biofilm growth reaching susceptibility after latest two days. For levofloxacin and piperacillin/tazobactam that were tested effective in CF-PA75, increase in MICs after one day or two days of biofilm growth, respectively, were observed reaching resistance levels (breakpoint 2 mg/L) for several colonies. However, we cannot explain this observation.

Most alarming was the strong increase in MICs of colistin already after one day of biofilm formation. The breakpoint of colistin is set at 4 mg/L. Most colonies recovered from biofilm showed MICs of 16 mg/L independently of the biofilm ‘age’.

We hypothesize that the observed changes in the susceptibility to the antibiotics is related to the mechanism of phage resistance as resistance to the virulent phage in CF-PA75 biofilm population occurred simultaneously after the first day of biofilm formation. In our recent study (ref. upon acceptance!), in which biofilms of various laboratory and clinical P. aeruginosa strains were treated sequentially with NP3, NP1, or M32 phages, we identified several mutations in structural genes of the type IV pilus machinery frequently present in clones resistant to these phages. It is unclear how these mutations are related to the restored susceptibility to different classes of antibiotics, since type IV pili are primarily known for adherence, motility, and uptake of free DNA (30–32)Another possibility involves changes in the lipopolysaccharides (LPS). In the previous study, we also linked some mutations in the LPS synthesis apparatus to phage resistance. Such changes would be consistent with the increased MIC values for colistin, as colistin is a surfactant and exerts its effects primarily through interaction with bacterial membranes (33). This alteration might reduce the ability of colistin to destabilise and permeabilize the LPs. It is however also unclear how this can reduce efficacy of β-lactams, fluroquinolones and aminoglycosides.

Prophages are known to contribute to the spread of antimicrobial resistance (AMR) and virulence factors. In P. aeruginosa, 12% of AMR genes and 10.5% of virulence factors were found to be associated with prophages within the bacterial genome. Notably, 1.1% of instances involved the co-location of both AMR genes and virulence factors. In Pf1-like prophages, only the *catB7* gene, which encodes a chloramphenicol acetyltransferase (CAT) and confers resistance to chloramphenicol, has been identified. This gene is not relevant for therapy. However, other prophages were found to contain genes conferring aminoglycoside resistance, such as *aph*(3’) and *ant*(3’), which could significantly impact CF therapy. It would therefore be interesting to determine whether these specific prophages (JBD25 and F10), classified as *Caudoviricetes* (head-tail structure), are also capable of lysogenic conversion and what effect this has on the resistance profiles of CF P. aeruginosa isolates.

## Materials and methods

### Bacterial strains

A total of 51 *P. aeruginosa* isolates from CF patients were examined retrospectively in this study (see also Supplementary Material Table S1). These originated from sputum, nasal lavage, and throat swap of CF patients collected at the Cystic Fibrosis Center for Children and Adults (Jena University Hospital, Germany) during the regular check-ups or acute symptoms of the patients. The *P. aeruginosa* strains were routinely isolated and confirmed by matrix assisted laser desorption ionization time-of-flight (MALDI-TOF) (Brucker Daltonics, Billerica, US) and the resistograms were performed using VITEK®2 (bioMerieux, Marcy l’ Etoile, France) at the Institute of Medical Microbiology (Jena University Hospital, Germany). The retrospective use of the isolates and selected pseudonymized data of the patients, was approved by the ethic committee of the Jena University Hospital (Germany) under the registration number 2023-3199-Material.

Laboratory *P. aeruginosa* strain PAO1 was used as control in several experiments, and to proliferate virulent phages and titrate the temperate and virulent phages,

### Phages

Three different virulent exogenous phages were used: NP1 phage (family *Siphoviridae*, *Nipunavirus*, GenBank: KX129925.1) with lytic activity against PA14 but not against PAO1 as well as NP3 phage (family *Myoviridae, Pbunavirus*, GenBank: KU198331.1), showing a broad range lytic activity against different *P. aueruginosa* strains, were initially isolated by Chaudhry *et al*., from a local sewage treatment plant (23); M32 phage (family *Autographiviridae,* Phikmvvirus, *G*enBank: KX711710.1) with lytic activity against PAO1 but not PA14, was isolated by Karumidze *et al*. from a Georgian sewage water (24).

### Culture conditions of bacteria

The *P. aeruginosa* strains were cultured on Columbia blood agar or cetrimide agar (all bioMerieux). Single colonies were cultured overnight (approximately 16 h) in a 4 mL Luria-Bertani (LB) broth (Carl Roth GmbH, Karlsruhe, Germany) at 37 °C with agitation of 160 rounds per minute (rpm) on an orbital shaker GFL 3032 (GFL Gesellschaft für Labortechnik GmbH, Burgwedel, Germany). These overnight cultures were used to prepare the experiments and cryo-stocks with 10% glycerol (V/V), which were stored at −80 ℃.

To grow biofilm, the overnight cultures were adjusted to an optical density (OD_600_) of 0.08 and grown in 96-well titre plates (Sarstedt AG &Co.KG, Nürnbrecht, Germany) in LB broth for several days (see specific methods below) under static conditions at 37°C. To determine viable bacteria in the biofilms, the biofilms were washed twice carefully with 0.9 % NaCl and the bacteria were resolved in fresh LB broth by scrapping the biofilms from the flat bottom 96-well titre plates. The bacteria suspension was serially diluted and 100 µl of selected dilutions were spread on LB agar, and grown overnight at 37°C.

### Phage proliferation and titration

Phage proliferation was performed using PA14 as the host for NP1 and NP3, and PAO1 as the host for M32. The bacteria were grown in LB broth until log-phase (OD_600_ 0.5 −0.8), adjusted to OD_600_ of 0.1 and 0.3 mL were mixed with 0.1 mL of phage lysate suspension and incubated for 30 minutes at 37 °C. Subsequently, 4 mL of LB top agar (0.5%) was added, and the phage-bacteria was overlayed on the LB-agar palate (1.2%) and incubated overnight at 37°C. To recover the phages, 5 mL of LB broth was poured over a plate with confluent but separate plaques and incubated for 2 hours with shaking at 80 rpm. The medium was transferred to 1.5 ml tubes and centrifuged. To obtain crude phage lysate, the supernatants were combined and filtered through a 0.2 µm syringe filter (Merck KGaA, Darmstadt, Germany) and stored at 4°C. To determine the phage titre, the same procedure was applied, but the phage lysates were diluted logarithmically before mixing with bacteria, and the plaques were counted after incubation.

### Isolation of induced prophage in biofilms

Strain PAO1 and clinical isolates CF-AP6, CF-PA75 and CF-PA83 were grown in LB broth without shaking for up to 6 days to form biofilms and supernatants were collected daily and fresh medium was added to the biofilms. The supernatants were filtered and treated by DNase I (1.25 U) (ThermoFischer Scientific) for 1 h at 37°C to remove free DNA. Subsequently the DNase was inactivated 75°C for 10 min.

To precipitate the phages in the crude phage lysate obtained from different wells (technical repeats), the lysate was subjected to a treatment with 10 % v/v PEG 6000 and 2.1 M NaCl (final concentrations) at 4°C overnight. The mixture was centrifuged for 30 min at 15000 rpm at 4°C in the centrifuge Eppendorf 5702RH (Eppendorf SE, Hamburg, Germany). The pellet was resuspended in the 1/5 of the in initial volumes in PBS buffer (ThermoFischer Scientific) and diluted with LB broth (1:1).

### Phage spot test and cross-streak agar assay

This test was used to qualitatively determine lytic activity in the supernatants of the induced prophages on its host strain (self-infectivity) or another bacterial strains (cross-infectivity). A fresh bacteria culture was adjusted to an OD_600_ of 0.1 and 0.3 mL of the bacteria suspension was mixed with 4 mL of top agar and poured onto an LB plate. After solidification, 10 µL of the lysate was spotted onto the plate and incubated overnight. The presence of clear plaques/halos indicated the susceptibility.

### Phage cross-streak agar assay

This test was used to qualitatively determine lytic activity of virulent phages. The phage lysates were adjusted to approximately 10^8^ PFU/mL and 40 µL were applied each as a line in the middle of a petri dish on LB agar plates. Before complete drying of the phage suspension, an inoculation loop of an overnight bacterial culture was streaked across the phage line. Per plate several bacterial cultured (up to six) were tested against one phage. The plates were incubated overnight at 37 ℃ and evaluated the next day. A missing growth after the phage line was interpreted as susceptibility and a coherent growth as resistance. Isolated colonies without uniform growth were considered as ‘reduced susceptibility’ (example see Supplementary Material Figure S1).

### DNA isolation of the strains

Genomic DNA was isolated from the bacterial pellet obtained from 1 ml overnight LB-cultre. The ZymoBIOMICS™ DNA Microprep kit (Zymo Research Corporation, Irvine, US) was uses according to the manufacturer’s protocol and quantity and purity of genomic DNA were assessed spectrometrically using Qubit fluorometer and the Qubit® dsDNA HS Assay (both ThermoFischer Scientific).

### Transmission of Pf4 in mixed biofilms

Overnight cultures of CF-PA75 and CF-PA6 were adjusted to OD_600_ of 0.08, mixed to equal volumes in 96-well titre plates (Sarstedt AG &Co.KG), and grown under static conditions at 37°C up to 5 days to form a mixed biofilm. The viable bacteria were assessed daily as described previously by selection for CF-PA75 and for CF-PA6 on LB ager supplemented with meropenem (8 g/L to select) or tobramycin (6 mg/L to select), respectively. From each strain and time point, five randomly selected colonies were taken, resolved in 100 µl sterile water and boiled for 10 min. After centrifugation for 10 min at 10,000 g and room temperature (RT), the presence of the Pf4 phage was assessed in the supernatants by specific PCR.

### Amplification of Pf1-like prophages elements

To determine prophage related genetic elements, previously described primers (each 200 nM) listed in Supplementary Material Table S2 (18,19), 100 ng of the genomic bacterial DNA and the Dream Taq DNA polymerase kit (ThermoFischer Scientific) were mixed according to the manufacturer’s recommendation in 20 µl volume. The PCR was performed in the Veriti 96 well thermo-cycler (Applied Biosystems, Waltham, USA) under the following conditions: an initial denaturation steps of 95 °C for 2 min, followed by 40 cycles of denaturation (95 ℃, 3 s), annealing (62 ℃, 2 s) extension (72 ℃, 1 min). DNA Gel Loading Dye (6X) was used to prepare the samples and O’GeneRuler DNA ladder was used as marker (both ThermoFischer Scientific). The amplicons were separated by electrophoresis on 2% agarose gels supplemented with ethidium bromide (0.2 μg/mL) at 100 V and documented using G-Box F3 (Syngene, Cambridge, UK).

### Antibiotic resistance assay via VITEK 2

We investigated the resistance pattern of randomly selected clones (each strain and treatment N = 32) isolated from biofilm after prophage induction for their susceptibility against several antibiotics. Individual wells were streaked out on an LB agar plate and incubated overnight. The selected colonies were resuspended each in 0.45% NaCl and adjusted to an OD600 0.50 – 0.63. The antimicrobial susceptibility profiles were assessed by VITEK®-2 system using the cartridges AST-N389 and AST-N232 for gram-negative organisms (all bioMérieux, Marcy-l’Étoile, France) according to the manufacture’s instruction.

### Software and statistical analysis

Microsoft Excel (Microsoft Corporation, Redmond, USA) and GraphPad Prism 9 (GraphPad Software Inc., San Diego, USA) were used for calculations, analyses and visualisation of experimental data.

## Acknowledgements

We gratefully acknowledge the technical assistance of Lisa Jasef and Lucy Buschack.

## Supplementary Material

Supplementary Metarial.docx

**Table S1** Patient cohort used in this study

**Table S2** Prophage primers used in this study

**Figure S1** Exemplary test of cross-streak agar assay

**Figure S2** Range of the patients’ age included in this study.

**Figure S3** Distribution of the antimicrobial susceptibility to antibiotics relevant in treatment of infections in CF- patients

**Figure S5.** Phage detection in the supernatants of biofilms of selected strains

